# eIF5B gates the transition from translation initiation to elongation

**DOI:** 10.1101/587022

**Authors:** Jinfan Wang, Alex G. Johnson, Christopher P. Lapointe, Junhong Choi, Arjun Prabhakar, Dong-Hua Chen, Alexey N. Petrov, Joseph D. Puglisi

## Abstract

Translation initiation determines both the quantity and identity of the protein product by establishing the reading frame for protein synthesis. In eukaryotic cells, numerous translation initiation factors (eIFs) prepare ribosomes for polypeptide elongation, yet the underlying dynamics of this process remain enigmatic^1–4^. A central question is how eukaryotic ribosomes transition from translation initiation to elongation. Here, we applied *in vitro* single-molecule fluorescence microscopy approaches to monitor directly in real time the pathways of late translation initiation and the transition to elongation using a purified yeast *Saccharomyces cerevisiae* translation system. This transition was remarkably slower in our eukaryotic system than that reported for *Escherichia coli*^5–7^. The slow entry to elongation was defined by a long residence time of eIF5B on the 80S ribosome after joining of individual ribosomal subunits, which is catalyzed by this universally conserved initiation factor. Inhibition of eIF5B GTPase activity following subunit joining prevented eIF5B dissociation from the 80S complex, thereby preventing elongation. Our findings illustrate how eIF5B dissociation serves as a kinetic checkpoint for the transition from initiation to elongation, and its release may be governed by a conformation of the ribosome complex that triggers GTP hydrolysis.

---

The assembly of an elongation-competent 80S ribosome complex requires choreographed interplay among numerous eIFs, the small (40S) and large (60S) ribosomal subunits, initiator Met-tRNAi, and mRNA^1–4^. In complex with eIFs and Met-tRNAi, the 40S subunit is recruited to an mRNA with a 7-methylguanosine (m^7^G) cap on its 5’ end and subsequently scans the 5’ untranslated region (5’UTR) to identify the correct AUG start codon (termed “scanning”). Upon AUG recognition, a series of compositional and conformational rearrangements occur, resulting in a 48S pre-initiation complex (PIC) that is primed for the 60S subunit to join and form the 80S complex^8–11^. The assembled 80S ribosome then transitions to the polypeptide-elongation phase of translation, marked by binding of an elongator aminoacyl-tRNA (aa-tRNA) to the ribosomal aa-tRNA binding site (A site) via a ternary complex with eukaryotic translation elongation factor 1A (eEF1A) and GTP^3^.

Despite decades of study, the transition from eukaryotic translation initiation to elongation remains poorly understood. Joining of the 60S subunit to the 48S PIC is catalyzed by the universally conserved factor eIF5B, independent of its GTPase activity^12–15^. The conformation of yeast eIF5B when bound to GTP is remodeled upon its association with a ribosome, enabling eIF5B to directly contact both Met-tRNAi and the GTPase activation center on the 60S subunit^16–18^. These contacts are likely essential for triggering GTP hydrolysis and the subsequent release of eIF5B-GDP, allowing the ribosome to enter the elongation phase^18^. Bypassing eIF5B GTP hydrolysis with a mutation that weakens its ribosome binding affinity reduces the stringency of start codon selection *in vivo*^14^, implying that the timing of eIF5B dissociation from the 80S complex and the subsequent entry into elongation is critical. The steps and their timescales that govern this transition have not been elucidated in eukaryotes because its complex, multi-step nature challenges traditional biochemical approaches. Here, we overcome these limitations by directly observing the process using single-molecule fluorescence microscopy to parse out the dynamics and pathways involved in 80S assembly and subsequent acceptance of the first elongator aa-tRNA.

We first reconstituted an active yeast translation system from purified ribosomes and core eIFs and eEFs (**Extended Data Fig. 1**) suitable for single-molecule analyses^19,20^. Ribosomal subunits were site-specifically labeled^21,22^ with donor and receptor fluorescent probes for unambiguous detection of 80S complex assembly by inter-subunit single-molecule Förster resonance energy transfer (smFRET) (**Fig. 1**). We tested our translation system using a previously well-characterized, uncapped, unstructured model mRNA (**Fig. 2a**) that functions independently of the cap-binding eIF4F complex, eIF4B and eIF3 for initiation^19,20^. Native gel shift experiments^20^ demonstrated that the components together could promote efficient 48S PIC and 80S formation on the model mRNA (**Extended Data Fig. 2a**). We next formed 80S complexes with Cy3-40S and Cy5-60S on the model mRNA to characterize the ribosomal smFRET pair with total internal reflection fluorescence microscopy (TIRFM) imaging, and the observed FRET efficiency was consistent with the estimated distance between the two labeling sites (**Fig. 1b,c,d, Extended Data Fig. 1d**). We then sought to extend our system to study real-time 80S assembly with a zero-mode waveguide (ZMW)-based instrumentation platform^23^, which is capable of multi-color single-molecule fluorescence imaging at high fluorescently-labeled ligand concentrations.

**Fig. 1.**
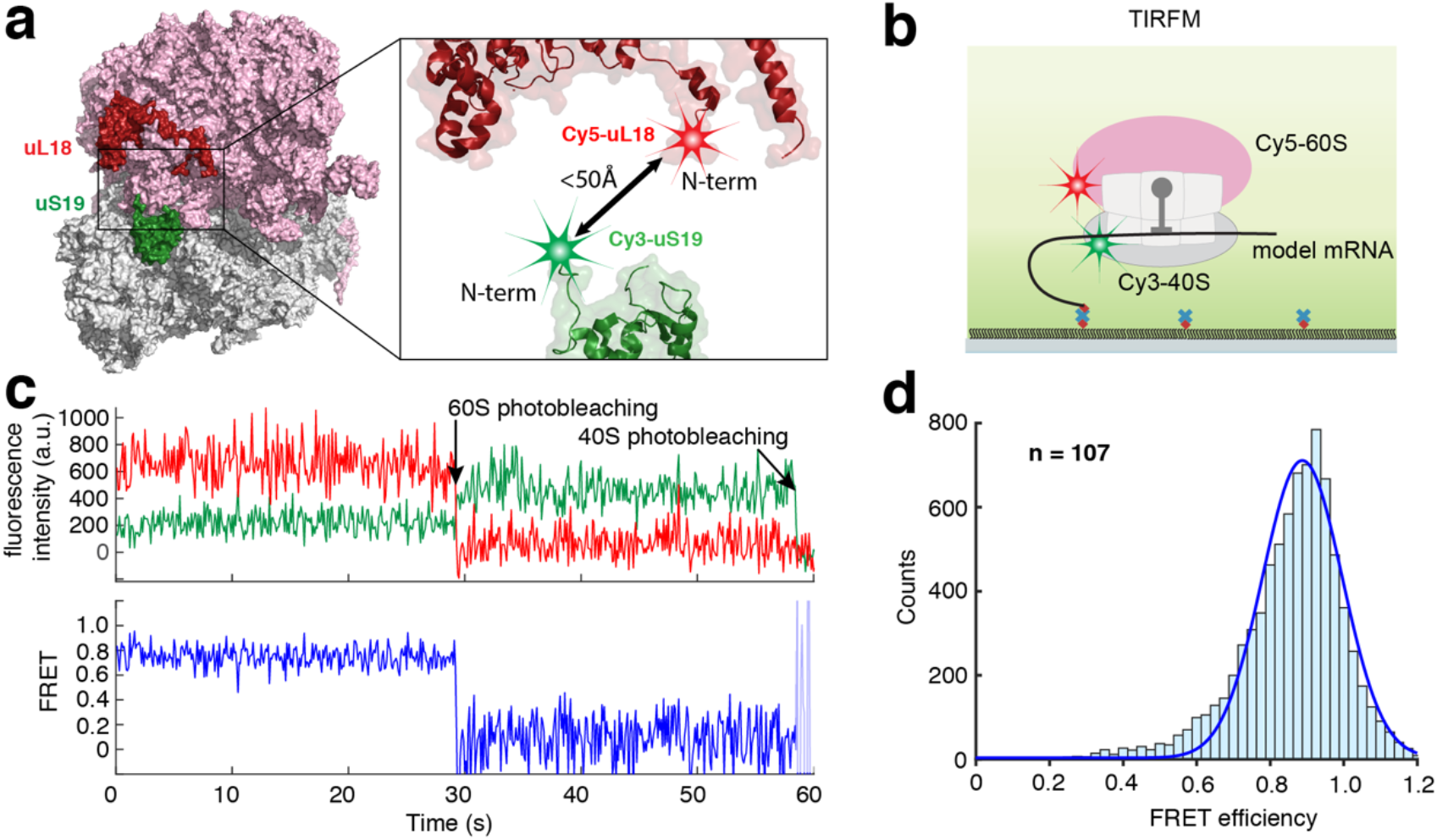
Engineering yeast ribosomes for inter-subunit smFRET. **a**, Yeast 40S ribosomal subunits were fluorescently labeled with a Cy3-like dye (referred to as Cy3) via a ybbR-tag^22^ at the N-terminus (N-term) of the ribosomal protein uS19, and the 60S subunits were labeled with a Cy5-like dye (referred to as Cy5) by a SNAP tag at the N-term region of uL18^21^. The estimated distance between the two labeling sites is within 50Å from published yeast 80S 3D structures (**Extended Data Fig. 1d**). The ribosome model was created in PyMOL with PDB 4V8Z^18^. **b**, TIRFM experimental setup to characterize the inter-subunit FRET signal. 80S complexes were assembled from Cy3-40S and Cy5-60S on the model mRNA (**Fig. 2a**) in the presence of required factors and were immobilized on a quartz slide used for TIRFM imaging with green laser illumination. **c**, Sample TIRFM experimental trace showing FRET conversions. **d**, The inter-subunit smFRET efficiency histogram was fit with a single-Gaussian distribution, with a mean FRET efficiency at 0.89 ± 0.15. Number of molecules analyzed (n) = 107.

**Fig. 2.**
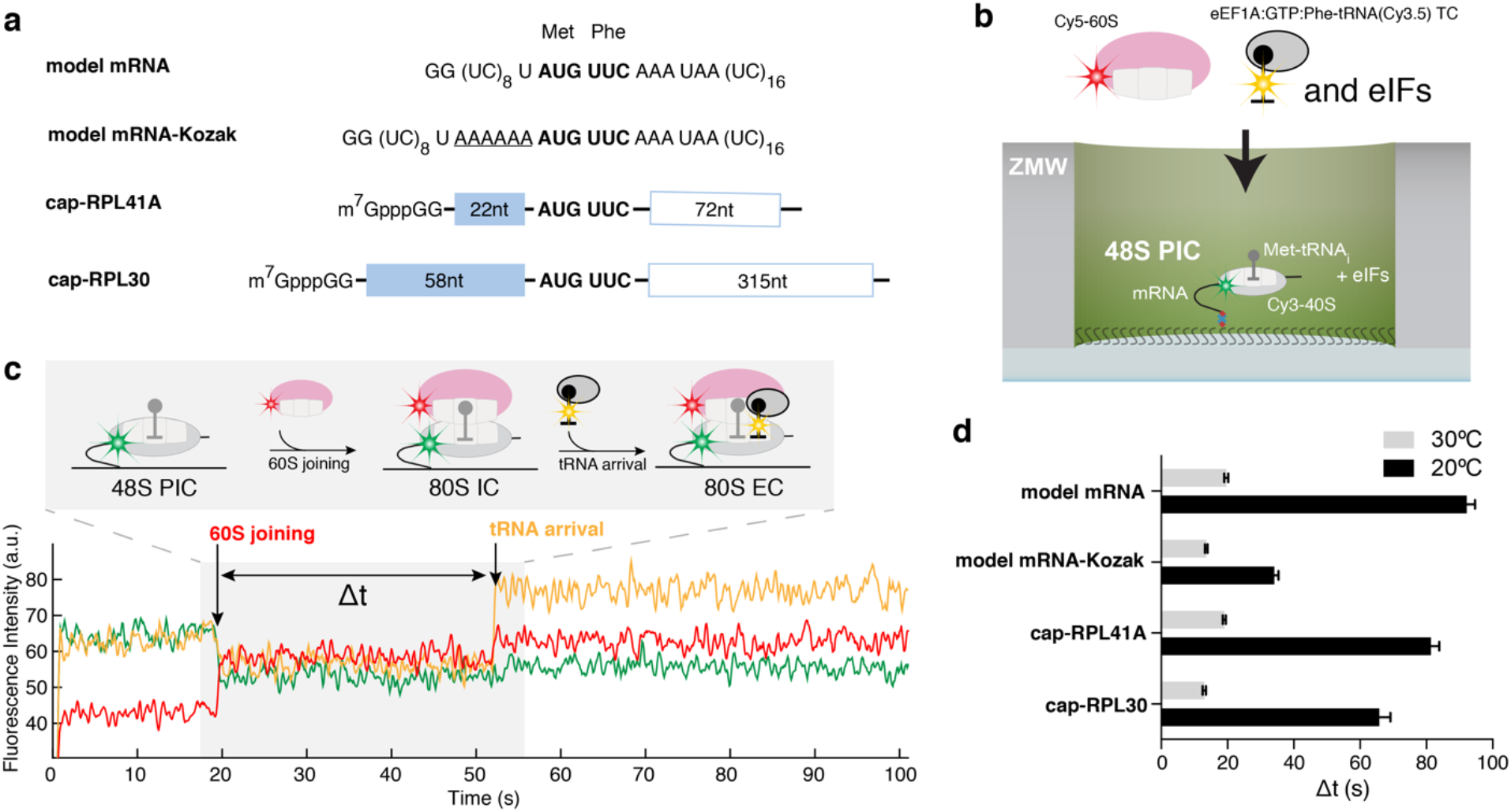
Real-time observation of eukaryotic translation initiation and the transition to elongation. **a**, mRNA constructs used in single-molecule assays. All the mRNAs contain a UUC phenylalanine (Phe) codon after the AUG start codon and are biotinylated at their 3’ ends. **b**, Experimental setup for single-molecule assays. 48S preinitiation complexes (PICs) containing Cy3-40S, Met-tRNAi, and the 3’-biotyinlated mRNA of interest were immobilized in ZMWs in the presence of required eIFs. Experiments were started by illuminating ZMWs with a green laser and delivering Cy5-60S, Cy3.5-Phe-tRNA^Phe^:eEF1A:GTP ternary complex (TC) and eIFs. **c**, Example experimental trace (bottom) and schematic illustration (top) of the molecular events along the reaction coordinate. In the example trace, 60S joining to the 48S PIC to form the 80S initiation complex (80S IC) was identified as the appearance of smFRET. The first A-site elongator tRNA binding was identified as a burst of yellow fluorescence, marking the formation of the 80S elongation complex (80S EC). The dwell times between the 60S joining and the A-site Phe-TC arrival were measured for many molecules, and the average of which, designated as the “Δt”, defines the average time for the transition from initiation to elongation. **d**, The Δt values compared across all assayed mRNAs and at 20°C and 30°C. Error bars represent the 95% confidence interval from fitting Δt values to single-exponential distributions. From bottom to top for each bar, number of molecules analyzed were 118, 130, 130, 121, 189, 159, 164 and 136.

Since we observed efficient and stable 48S PIC formation on the mRNA in native gel shift experiments (**Extended Data Fig. 2a**), we formed and immobilized single 48S PICs (with Cy3-40S) within ZMWs and delivered Cy5-60S along with the other required factors (**Extended Data Fig. 3a**). We observed efficient 60S subunit joining on single 48S PICs, with ~ 90% of the 48S PICs binding to Cy5-60S to form the 80S complex (**Extended Data Fig. 3b**), which required eIF5B; in the absence of eIF5B, no 80S formation was observed in the 15-min imaging time window (n = 500). The association rate of 60S subunits was comparable to that from prior bulk measurements (**Extended Data Fig. 3c**)^15^. When the first elongator Phe-(Cy3.5)-tRNA^Phe^ was included in the delivery mix as a ternary complex with eEF1A and GTP (Cy3.5-Phe-TC) (**Fig. 2b**), ~80% of the 60S joining smFRET events were followed by a subsequent Cy3.5-Phe-TC arrival event (**Fig. 2c**). These results demonstrated that real-time 80S assembly occurred in the correct reading frame and that our single-molecule translation initiation system was fully active.

To determine an initial rate for the transition from initiation to elongation, we measured the dwell times between 60S joining and the arrival of the Cy3.5-Phe-TC and determined the average time of this transition (“Δt”) (**Fig. 2c**). The Δt was 92.2 ± 2.5 sec when assayed at 20°C in the same buffer (containing 3 mM free Mg^2+^) as used in the bulk kinetic analyses of yeast translation initiation (**Fig. 2d**)^15^. This was surprisingly longer than the bacterial counterpart, which was reported to be ~1 sec at 20°C^5–7^. The Δt remained similar upon addition of 150 nM eIF3 and 200 nM eEF3 (and 1 mM ATP:Mg^2+^), or 500 nM fully hypusine-modified eIF5A^24^, indicating that the transition from initiation to elongation is independent of these factors (**Extended Data Fig. 4a**). Decreasing the concentration of Cy3.5-Phe-TC did not significantly alter the Δt (**Extended Data Fig. 4b**). Thus, the rate of the transition was limited by a step prior to binding of the first A-site tRNA.

We next tested whether the slow transition was due to the cap-independence of initiation on the model mRNA (*i.e*., the missing m^7^G cap on the model mRNA and eIF4F proteins in the reaction). We formed 48S PICs on cap-RPL41A or cap-RPL30 mRNAs (**Fig. 2a**, mRNAs were completely capped with m^7^G, see Online Methods) in the presence of eIF4F, eIF4B and eIF3 proteins and performed the same experiments as above. The 48S PICs readily formed in a cap-dependent manner (**Extended Data Fig. 2b,c**) and were in the post-scanning, mRNA channel-“closed” state as demonstrated by cryo-electron microscopy analyses (**Extended Data Fig. 2d,e,f**)^11^. Thus, the 48S PICs central to our single-molecule assays were an authentic and on-pathway complex. Yet, the Δt values were 81.4 ± 2.5 sec and 65.7 ± 3.4 sec for cap-RPL41A and cap-RPL30, respectively, which were similar to the Δt measured on the model mRNA (**Fig. 2d**). Increasing the Cy3.5-Phe-TC concentration did not change the Δt value (**Extended Data Fig. 4c**), further demonstrating that the rate-limiting step occurs prior to A-site tRNA binding. Our findings suggest that the slow transition to elongation may be a general feature of yeast translation.

We next tested whether the sequence context near the start codon or reaction temperature impacted entry into the elongation phase of translation. When the optimal yeast Kozak sequence^25,26^ was inserted just upstream of the AUG codon on the model mRNA (designated as model mRNA-Kozak), the Δt was reduced by ~3-fold to 34.1 ± 1.3 sec (**Fig. 2a,d**). Thus, while the transition rate was altered by the sequence context surrounding the start codon, even the fastest rate we observed was still more than 30-fold slower than that observed in *E. coli*^5–7^. To understand the energetics of the rate-limiting step in the eukaryotic transition, we performed the same experiments at an increased temperature (30°C), and found a striking dependence of Δt on temperature (**Fig. 2d**): a ~4-5-fold decrease for the model and cap-RPL41A/30 mRNAs, and a ~2.5-fold decrease for the model mRNA-Kozak.

The temperature dependence of Δt suggested that enthalpically-driven slow conformational rearrangements and/or factor dissociation limited the transition. We hypothesized that eIF5B dissociation was the rate-limiting step given that its binding site on the ribosome overlaps with that of the elongator aa-tRNAs^18^. If eIF5B indeed gates the transition to elongation, we expect that its lifetime on the 80S would be similar to the Δt measured above.

To test this hypothesis, we fluorescently labeled eIF5B (**Fig. 3a**) and tracked the protein throughout the reaction pathways using a similar experimental scheme as above. A Cy5.5 dye was attached to N-terminally truncated eIF5B (remaining residues 396-1002), via a ybbR-tag fused to its N-terminus. This truncated eIF5B is the standard construct in the eukaryotic translation field^12,15,27,28^ and was used in all our measurements above (**Extended Data Fig. 5a**), and neither the tag nor dye impacted eIF5B function (**Extended Data Fig. 5**). After immobilizing pre-formed Cy3-48S PICs in ZMWs, we co-delivered Cy5.5-eIF5B, Cy5-60S and Cy3.5-Phe-TC in the presence of other required factors to the immobilized complex. With the model mRNA and direct illumination of all fluorophores, we observed transient Cy5.5-eIF5B sampling events to the 48S PIC prior to 60S joining (**Extended Data Fig. 6a**), consistent with its dynamic interaction with that complex as suggested by ensemble fluorescence measurements^15^. Upon Cy5-60S joining to form the 80S complex, as signaled by smFRET, the average Cy5.5-eIF5B lifetime on the ribosome was prolonged and nearly equal to Δt (**Fig. 3b,c,d, Extended Data Fig. 6a**). Importantly, the Δt measured here matched that measured with unlabeled eIF5B, reassuring us of the non-perturbed function post labeling (**Extended Data Fig. 5d**). Furthermore, Cy5.5-eIF5B dissociation always preceded Cy3.5-Phe-TC binding to the A-site of the 80S ribosome, which occurred very rapidly (within 1 sec) after eIF5B departure (**Fig. 3b,e**). Similar results were observed with the other mRNAs at both 20°C and 30°C (**Fig. 3d**). Notably, the Δt was not reduced by using a full-length eIF5B, indicating that the lack of the non-conserved N-terminal region^14,16^ of the protein did not interfere with the transition rate (**Extended Data Fig. 5f**). Thus, the presence of eIF5B on the *de novo* assembled 80S ribosome limits the rate of the transition from initiation to elongation.

**Fig. 3.**
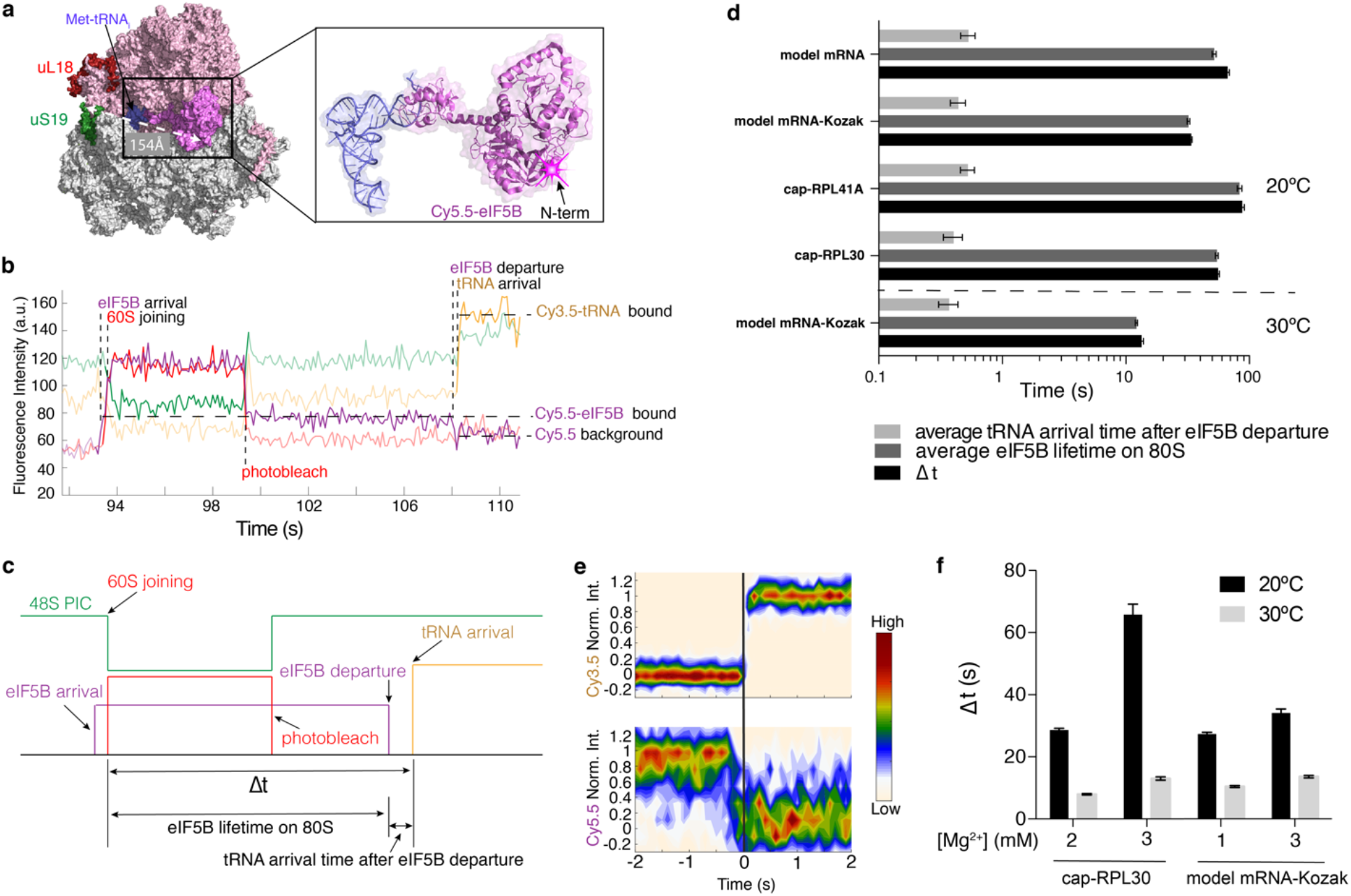
eIF5B gates the transition between initiation and elongation. **a**, Fluorescent labeling of eIF5B with Cy5.5 via a ybbR-tag at the N-terminal end, which is distal from the ribosomal subunit labels and hence not expected to interfere with the inter-subunit smFRET. The ribosome model was created in PyMOL with PDB 4V8Z^18^. **b**, Sample trace showing the correlation of the Cy5.5-eIF5B occupancy on the ribosomal complex with Cy5-60S joining and Cy3.5-Phe-TC arrival in experiments performed using the model mRNA. **c**, Schematic illustration of the molecular events in Fig. 3b. **d**, The Δt values, the average eIF5B lifetimes on 80S, and the average tRNA arrival times after eIF5B departure from experiments performed with Cy5.5-eIF5B compared across all assayed mRNAs. From bottom to top for each group of three bars, n = 141, 164, 133, 131 and 134. **e**, Contour plots of the Cy5.5-eIF5B departure (bottom, normalized Cy5.5 fluorescence intensity changing from 1 to 0) and Cy3.5-Phe-TC arrival (top, normalized Cy3.5 intensity changing from 0 to 1) in experiments performed with model mRNA in the presence of GTP, generated by superimposing all the analyzed fluorescence traces such that the Cy3.5-Phe-TC arrival was set at time 0. Contours are plotted from tan (lowest population) to red (highest population). Number of molecules analyzed (n) = 134. **f**, The Δt values from experiments performed with unlabeled eIF5B and cap-RPL30 or model mRNA-Kozak under denoted conditions. From left to right for each bar, n = 195, 152, 118, 130, 150, 132, 189 and 159. Error bars in **d** and **f** represent 95% confidence intervals from fitting the lifetimes to single-exponential distributions.

The rate of GTP hydrolysis by eIF5B is thought to define its residence time on the 80S complex. The Δt measured here on the model mRNA was similar to the mean time of eIF5B GTP hydrolysis during initiation on the same type of model mRNA^27^. Importantly, inclusion of the non-hydrolysable GTP analog GDPNP trapped eIF5B on the 80S complex (average Cy5.5-eIF5B lifetime on the 80S was ~850 sec and photobleaching/imaging-time limited, n = 105), and prevented A-site aa-tRNA association (**Extended Data Fig. 6b,c**). Thus, eIF5B departure is indeed controlled by GTP hydrolysis. Since free Mg^2+^ concentration influences the conformation of the ribosome^29^ and eIF5B GTP hydrolysis is triggered by the 60S subunit, we measured the effect of free Mg^2+^ concentration on Δt at increasing temperatures. The rate of transition to elongation decreased (*i.e*., the Δt value increased) with increasing free Mg^2+^ concentration while at higher temperature the transition was faster (**Fig. 3f and Extended Data Fig. 7a**), and consistently, the changes in rates were due to altered eIF5B residence times on the 80S (**Extended Data Fig. 7b**).

The striking dependence of eIF5B departure on temperature and free Mg^2+^ concentration suggests that its GTP hydrolysis activity is sensitive to the conformation of the ribosome. This is consistent with proposals derived from structural analyses^16–18,30^. Upon subunit joining, the 80S ribosome likely adopts a conformation wherein the 40S subunit is in the fully rotated state in relation to the 60S subunit, which orients the catalytic-site histidine (H480) of eIF5B in an inactive configuration. Activation of eIF5B GTPase activity requires a partial back-rotation of the 40S subunit to reposition a universally conserved tyrosine residue (Y837) in eIF5B and stabilize H480 in the catalytically active conformation^30^. We suspect that varying the temperature and free Mg^2+^ concentration in our experiments altered the energy landscape of these conformational rearrangements and thus the rate of GTP hydrolysis before eIF5B departure. Furthermore, in the pre-GTP hydrolysis state, eIF5B makes extensive contacts around the essential discriminating A1-U72 base pair of the Met-tRNAi, coupling the initiator tRNA recognition to its GTPase activation by the ribosome^18,31^. A GTPase-defective eIF5B mutant (H505Y), which may bypass the requirement of GTP hydrolysis for its departure due to its weakened ribosome binding affinity, leads to inaccurate start codon selection in vivo^14^. The H505Y mutant eIF5B did not significantly alter the subunit joining rate in our assays (**Extended Data Fig. 8a**), which began post scanning. Instead, initiation with H505Y eIF5B showed an ~15-fold decrease of the Δt value compared with that of wild-type eIF5B, independently of GTP hydrolysis, thus reducing the eIF5B occupancy time on the assembled 80S complex (**Extended Data Fig. 8b**). Collectively, our findings indicate that the rate of GTP hydrolysis by eIF5B and its subsequent dissociation serve as the final kinetic checkpoint of translation initiation prior to entry into the energy-expensive elongation phase.

In summary, we demonstrate that eIF5B gates the transition from eukaryotic translation initiation to elongation (**Fig. 4**). The lowest transition time we measured in m^7^G-cap-dependent translation was ~8 sec in the presence of 2 mM free Mg^2+^ at 30°C (**Fig. 3f**) – a timescale relevant to the estimated rates of translation initiation *in vivo*^32,33^. We speculate that the long lifetime of eIF5B-bound 80S complexes could partially account for the enrichment of ribosomal protected mRNA fragments at the start codon observed in ribosome profiling experiments in eukaryotic systems^34–40^. The relatively slow transition between translation initiation and elongation in the eukaryotic system versus the bacterial counterpart would render a mechanism to control the ribosomal density in the early part of the mRNA open reading frame^32,37^ to prevent ribosome collisions^41,42^, and a kinetic checkpoint for further quality control and regulation. The relatively long residence time of eIF5B on the 80S initiation complex may also be an evolutionary compromise due to its parallel role during ribosome biogenesis wherein it catalyzes 60S joining to immature 40S subunits as a final quality control check^43,44^. Nevertheless, we expect that there are many kinetic checkpoints in the early stages of eukaryotic translation. The application of similar single-molecule approaches will unveil the complicated dynamics underlying mRNA selection, scanning and start codon selection, and their regulation^1–4^.

**Fig. 4.**
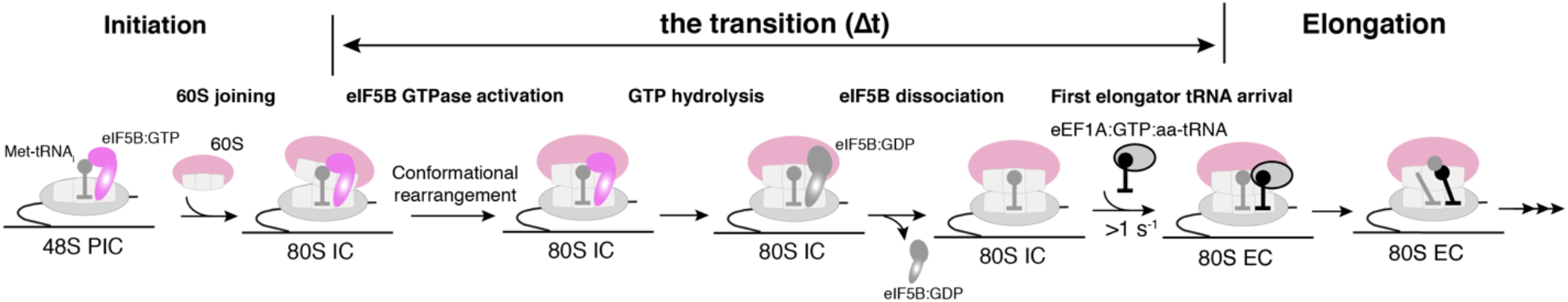
Model of the late eukaryotic translation initiation and its transition to elongation. eIF5B catalyzes 60S subunit joining to the 48S PIC to form the 80S initiation complex (IC), and its dissociation from the 80S IC requires GTP hydrolysis, plausibly leading to an altered eIF5B conformation thereby lowering its affinity to the 80S. Thus, the dissociation of eIF5B from the 80S IC gates the transition to elongation, marked by the binding of an elongator aa-tRNA to the elongation 80S complex (80S EC). Effects of free Mg^2+^ concentration, the sequence context surrounding the start codon, and temperature on the rate of the transition indicate that conformational rearrangements may play key roles in governing the rate of eIF5B dissociation, likely by controlling its GTPase activity.

## Acknowledgements

We are grateful to J. Lorsch, T. Dever, J. Dinman, J. Yin and C. Aitken for sharing constructs, strains and protocols; C. Sitron and O. Brandman’s lab for generously sharing equipment and knowledge for use of their Freezer/Mill for yeast lysis; the Stanford PAN facility for protein mass spectrometry analyses; and members of Puglisi laboratory for discussion and input. This work was supported by the US National Institutes of Health (NIH) grant GM011378 to J.D.P.; a Knut and Alice Wallenberg Foundation postdoctoral scholarship to J.W.; a National Science Foundation Graduate Research Fellowship (DGE-114747) to A.G.J.; a Damon Runyon Fellowship funded by the Damon Runyon Cancer Research Foundation (DRG-#2321-18) to C.P.L.; a Stanford Bio-X fellowship to J.C.; a Stanford Interdisciplinary Graduate Fellowship and NIH Molecular Biophysics Training Grant T32-GM008294 to A.P..

## Author Contributions

J.W. performed all the biochemical and single-molecule experiments. J.W. analyzed the data with the help from J.C., A.P., and A.N.P. D.-H.C. and J.W. acquired and D.-H.C. processed the cryo-EM data. J.W. and J.D.P. conceived the project with input from A.G.J., C.P.L and A.N.P. J.W., A.G.J., C.P.L., J.C., D.-H.C., and J.D.P. wrote the manuscript.

