## Extended Data Figures for "eIF5B gates the transition from translation initiation to elongation"

[5]Present address: Department of Biological Sciences, Auburn University, Auburn, AL 36849, USA

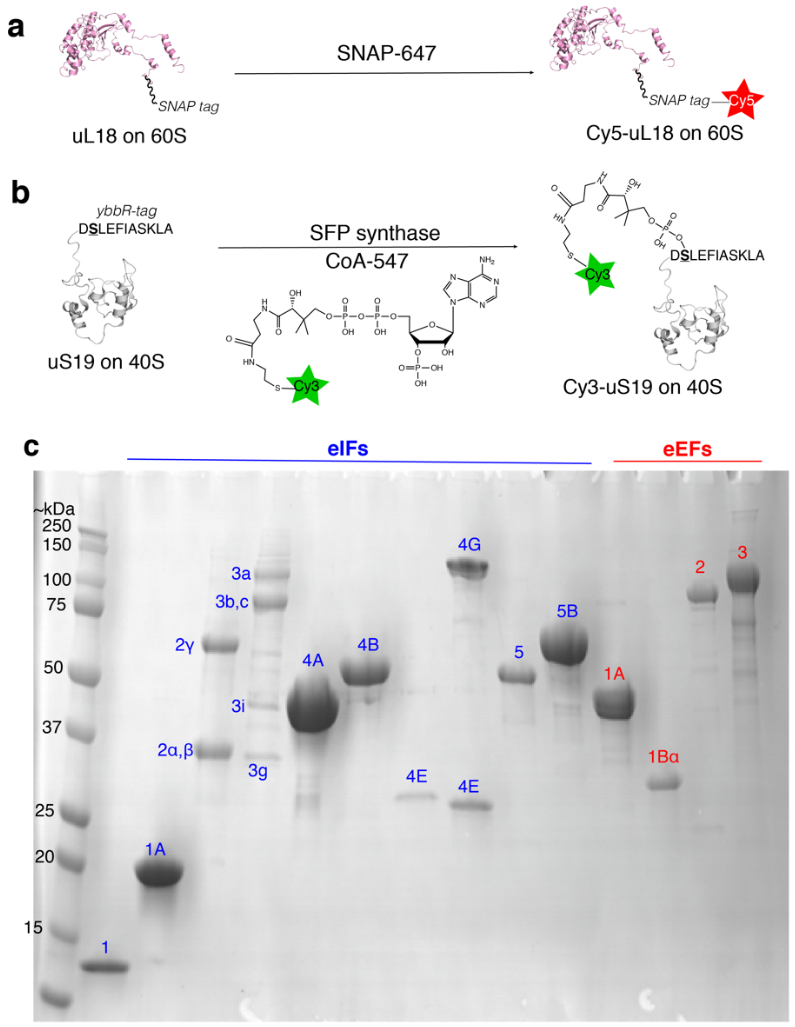


**d**

| PDB | brief description | Distance between the resolved N-terminal residues | | | expected FRET efficiency |
|---|---|---|---|---|---|
| | | uS19 | uL18 | distance (Å) | |
| 5JUO | eEF2-GDP-sordarin and TSV IRES bound, fully rotated | K8 | A2 | 45 | 0.74 |
| 3J77 | rotated ribosome bound with 1 tRNA | K8 | A2 | 40 | 0.87 |
| 5M1J | Dom34 and Hbs1 bound | K8 | A2 | 39 | 0.87 |
| 4V8Z | eIF5B and Met-$tRNA_i$ bound initiation complex | K8 | Q3 | 39 | 0.88 |
| 6GQ1 | mRNA, tRNA and eEF2 (GMPPCP/sordarin) bound | K8 | A2 | 39 | 0.88 |
| 5ON6 | haemanthamine bound | K8 | A2 | 38 | 0.89 |
| 4V88 | Stm1 bound | K8 | A2 | 38 | 0.89 |
| 5JUU | eEF2-GDP-sordarin and TSV IRES bound, least rotated | K8 | A2 | 35 | 0.93 |
| 3J78 | non-rotated ribosome bound with 2 tRNAs | K8 | A2 | 34 | 0.94 |
| 5MC6 | Ski2-Ski3-Ski8 complex bound | K8 | A2 | 32 | 0.96 |

**Extended Data Fig. 1. A reconstituted yeast translation system with fluorescently labeled ribosomes for inter-subunit smFRET.**

**a**, We previously have established the Cy5 labeling of yeast 60S ribosomal subunit via a SNAP tag fused to the uL18 protein and the reaction with a SNAP-647 dye[1].

**b**, In this work, we engineered a yeast strain in which all the 40S subunits carried the N-terminal ybbR-tagged uS19 protein. Upon purification, the 40S was labeled by SFP synthase with CoA-547 at the serine residue (in bold and underlined) of the ybbR tag, resulting in the Cy3-40S.

**c**, A representative SDS-PAGE analysis of the purified core eIFs (blue numbering) and eEFs (red numbering) used for the reconstitution of the translation system.

**d**, Estimated distances between the two labeling sites on the ribosomal subunits from a few examples of the published yeast 80S structures in different functional states, and the expected FRET efficiencies based on a Förster radius ($R_0$) of 54 Å for the Cy3/Cy5 FRET pair[2].

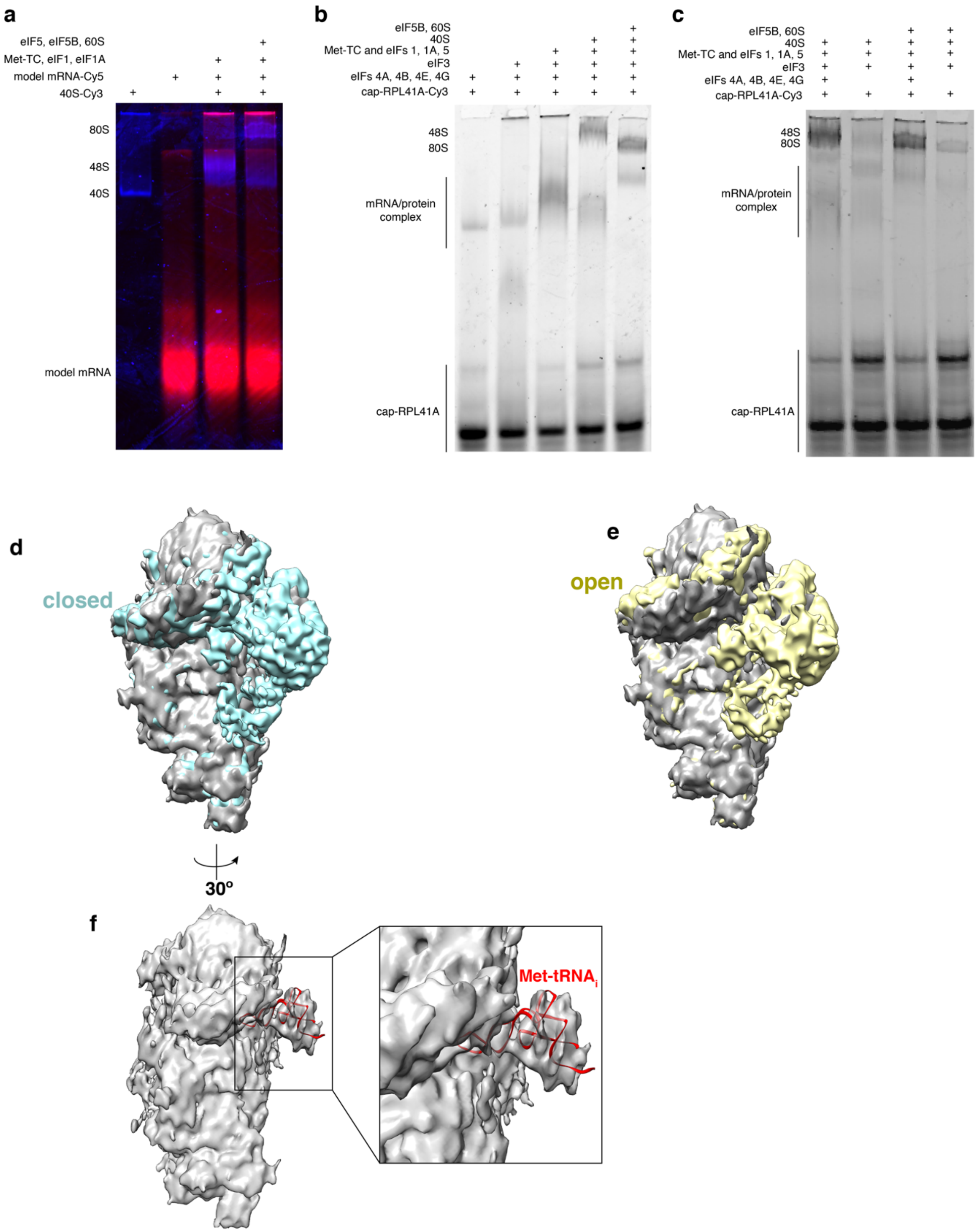


**Extended Data Fig. 2. Native gel shift assays and cryo-electron microscopy analysis showing active translation initiation with our purified yeast translation system.**

**a**, A representative gel showing initiation on the model mRNA. A merged view of Cy5 (red) and Cy3 (blue) scans of the same gel is shown. The model mRNA was labeled with Cy5 and 40S was labeled with Cy3. Addition of Cy3-40S, Met-tRNA$_i$:eIF2:GTP (Met-TC), eIF1 and eIF1A to the model mRNA-Cy5 resulted in the formation of a distinct 48S PIC band. Further addition of eIF5, eIF5B and 60S led to the formation of the 80S band.

**b**, A representative gel showing initiation on the cap-RPL41A mRNA. The cap-RPL41A mRNA was labeled with Cy3, and other components were unlabeled. The gel was scanned for Cy3 fluorescence. Various mRNA/protein

complexes were formed in the absence of the 40S. Upon adding 40S to the mixture, a distinct 48S PIC band was formed. The addition of eIF5B and 60S to the 48S PIC formation mixture resulted in the 80S band formation.
**c**, A representative gel showing initiation on the cap-RPL41A mRNA (fully $m^7G$-capped, see Online Methods) was via the cap-dependent pathway. Both 48S PIC and 80S formation were very inefficient when the cap-binding eIF4F (eIFs 4A, 4E and 4G) and eIF4B proteins were omitted from the reaction, demonstrating the cap-dependence of the initiation when the full set of eIFs were added.
**d**,**e** and **f**, With our 48S PIC assembly regime, we would expect that the 48S PIC would be in the post-scanning state, with the eIFs required during the scanning process potentially dissociated from the complex. A 9.9Å cryo-EM map was obtained for the 48S PIC formed on the cap-RPL30 mRNA (grey), which was compared with the reported scanning-incompetent, mRNA channel-closed (EMD 3048, cyan, **d**) or scanning-competent, mRNA channel-open (EMD 3049, yellow, **e**) 48S PIC structures[3] (see Online Methods). The cap-RPL30/48S PIC was assembled in the same way as for our single-molecule experiments, and the comparisons showed that it resembles the post-scanning closed state, with the Met-tRNA$_i$ positioned in the P-site (**f**, in red was the modeled Met-tRNA$_i$ in the EMD 3048 structure, PDB 3JAP).

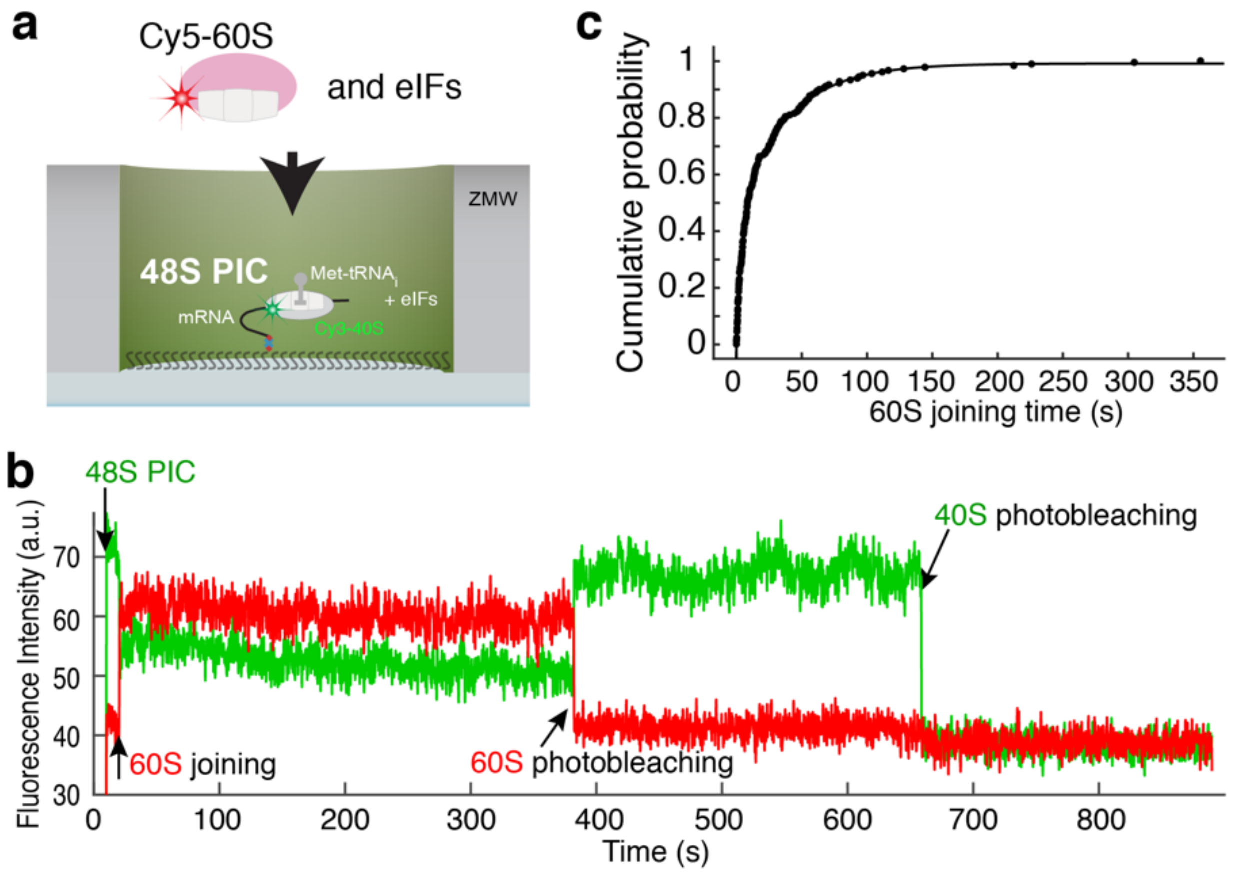


**Extended Data Fig. 3. Efficient real-time 80S assembly on the model mRNA at the single-molecule level.**
**a**, The smFRET assay for subunit joining in ZMWs. The 48S PICs were formed by incubating Cy3-40S, Met-TC, model mRNA-biotin, eIF1, eIF1A and eIF5 at 30ºC for 15 min before immobilization in the ZMWs. After washing away free components, the experiment was started with green laser illumination and delivery of Cy5-60S, eIF5, and eIF5B. The reaction was performed in the 1x Recon buffer supplemented with 1 mM GTP:$Mg^{2+}$ at 20ºC.
**b**, Example experimental trace showing real-time observation of Cy5-60S joining to immobilized Cy3-48S PIC to form the 80S complex, identified by the appearance of mFRET. Single photobleaching events are denoted.
**c**, Kinetics of 60S joining was fit to a double-exponential equation, resulting in a fast phase rate of ~0.22 $s^{-1}$ with ~46% amplitude, and a slow phase rate of ~0.03 $s^{-1}$ with ~54% amplitude (number of molecules analyzed n = 178). The kinetics is comparable to prior bulk measurement of the same reaction under similar condition (~77% fast phase with a rate of ~0.076 $s^{-1}$; ~23% slow phase with a rate of 0.019 $s^{-1}$)[4].

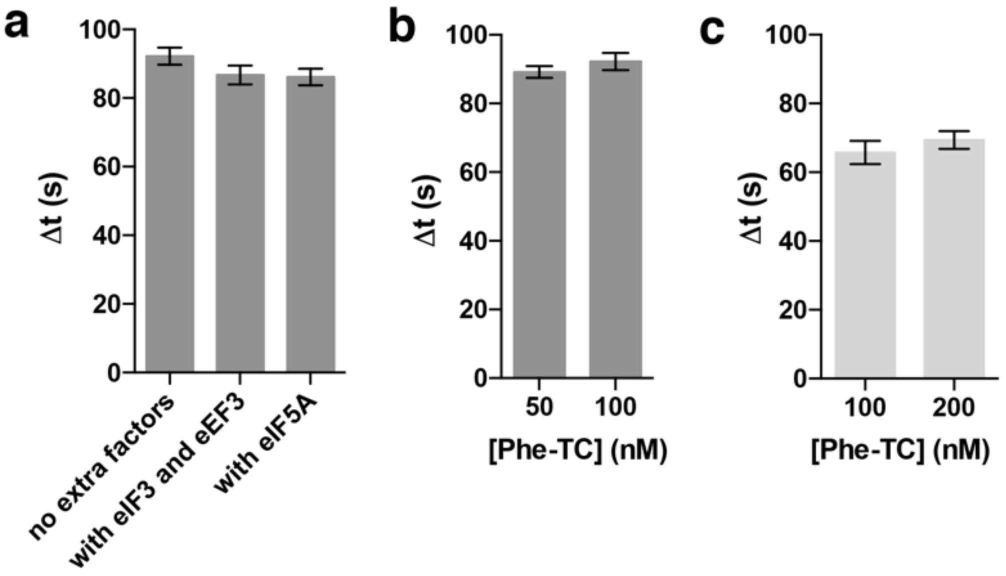


**Extended Data Fig. 4. Comparisons of the kinetics of the transition from initiation to elongation under various conditions.**

**a**, The transition kinetics were compared when no extra factors were added (data taken from **Fig.2d**, n = 164), or in presence of eIF3 and eEF3 (n = 130), or with addition of eIF5A (n = 143) in experiments performed with the model mRNA.

**b**, The transition kinetics were compared when the concentration of Cy3.5-Phe-TC was at 50 nM (n = 150) or 100 nM (data taken from **Fig.2d**, n = 164) in experiments performed with the model mRNA.

**c**, The transition kinetics were compared when the concentration of Cy3.5-Phe-TC was at 100 nM (n = 118) or 200 nM (n = 124) in experiments performed with the cap-RPL30 mRNA.

Error bars represent the 95% confidence intervals from fitting of the lifetimes to single-exponential distributions.

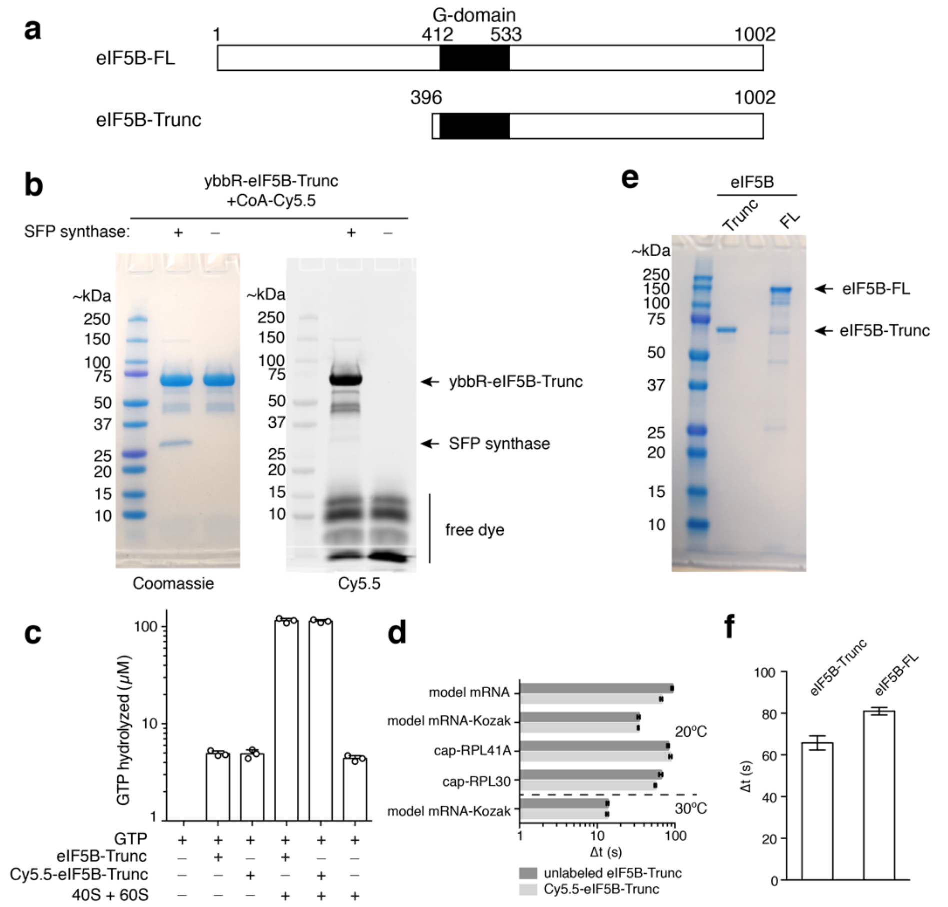


**Extended Data Fig. 5. Neither truncation nor fluorescent labeling of eIF5B perturbs its function.**
**a**, A N-terminal domain truncated version of eIF5B (eIF5B-Trunc) was used in most of our assays, as in other reported reconstituted, purified yeast translation assays[4–12]. Previous reports failed to purify the full-length protein and have demonstrated that the truncated protein supported initiation *in vitro* and *in vivo*[4,12].
**b**, We tagged eIF5B-Trunc at the N-terminus with a ybbR tag and labeled the protein by SFP synthase with a CoA-Cy5.5 dye. A representative gel is shown, which was first scanned for Cy5.5 fluorescence (right) and subsequently stained with Coomassie blue (left) following SDS-PAGE analysis of ybbR-eIF5B-Trunc post-labeling with and without SFP synthase.
**c**, The GTPase activity of Cy5.5-eIF5B-Trunc was not perturbed by the labeling. Multiple turnover GTP hydrolysis was performed in 50 mM HEPES-KOH pH 7.5, 10 mM $Mg(OAc)_2$, 100 mM KOAc at 30ºC 30min before quenching with malachite green assay solution. Where applicable, concentrations were: GTP 100 µM; eIF5B-Trunc 2.5 µM; Cy5.5-eIF5B-Trunc 2.5 µM; 40S+60S 0.2 µM each. The GTP only group was used as negative controls and the values were normalized to 0. Bars represent mean, and error bars indicate standard deviations of three replicates (individual data points are indicated with open circles).
**d**, The Δt values in experiments performed with Cy5.5-eIF5B (related to **Fig.3d**) versus those with unlabeled eIF5B (related to **Fig.2d**) compared across all assayed mRNAs and at 20ºC and 30ºC. From bottom to top for each bar, n = 141, 159, 164, 118, 133, 130, 131, 189, 134 and 164. Error bars represent the 95% confidence intervals from fitting of the Δt values to single-exponential distributions.
**e**, Despite it being reported that recombinant yeast eIF5B-FL purification cannot be achieved[13], we were able to recombinantly express and purify it as shown by a 12% SDS-PAGE gel analysis.
**f**, Use of the full-length eIF5B in our assay did not lead to faster transition to elongation. The Δt values were from experiments performed with the cap-RPL30 mRNA at 3 mM free $Mg^{2+}$ 20ºC. Error bars represent the 95% confidence intervals from fitting of the lifetimes to single-exponential distributions. For bars from left to right, n = 118 (related to **Fig.2d**) and 205.

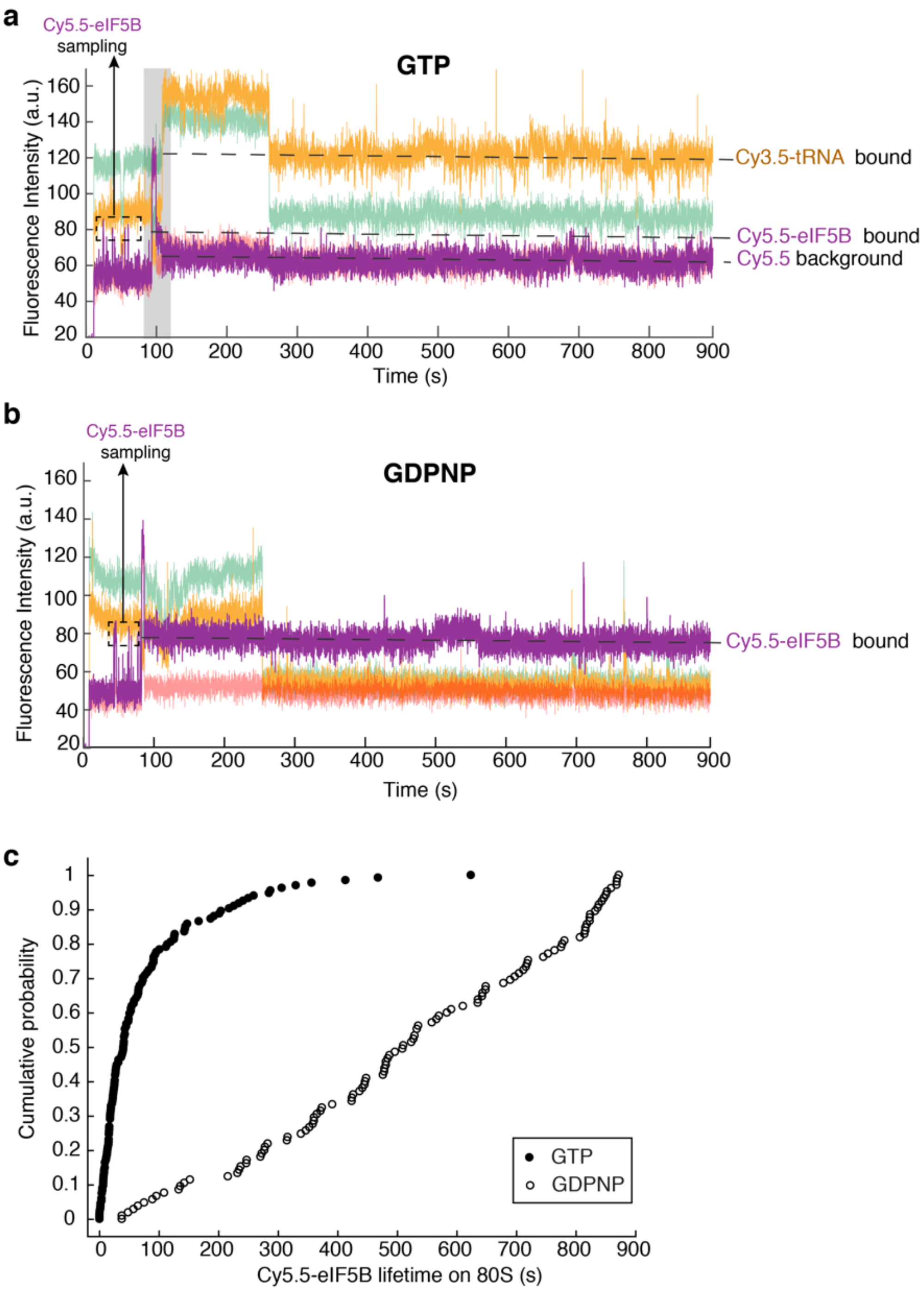


**Extended Data Fig. 6. The use of a non-hydrolysable GTP analog, GDPNP, traps eIF5B on the 80S and prevents the transition to elongation.**
**a**, Sample trace from experiments performed with GTP. The grey highlighted part of the trace is shown in **Fig. 3b** as a zoomed-in view.
**b**, Sample trace from experiments performed with non-hydrolysable GDPNP, whereupon 60S joining eIF5B is trapped on the 80S and no A-site tRNA binding was observed.
Black dash boxed Cy5.5 events are transient eIF5B sampling events to the 48S PIC prior to 60S joining.
**c**, The kinetic curves showing Cy5.5-eIF5B lifetime on 80S in experiments performed with the model mRNA at 3 mM $Mg^{2+}$ and 20ºC in the presence of GTP (51.9 ± 1.8 s, n = 134, related to **Fig. 3d**) or GDPNP (n = 105).

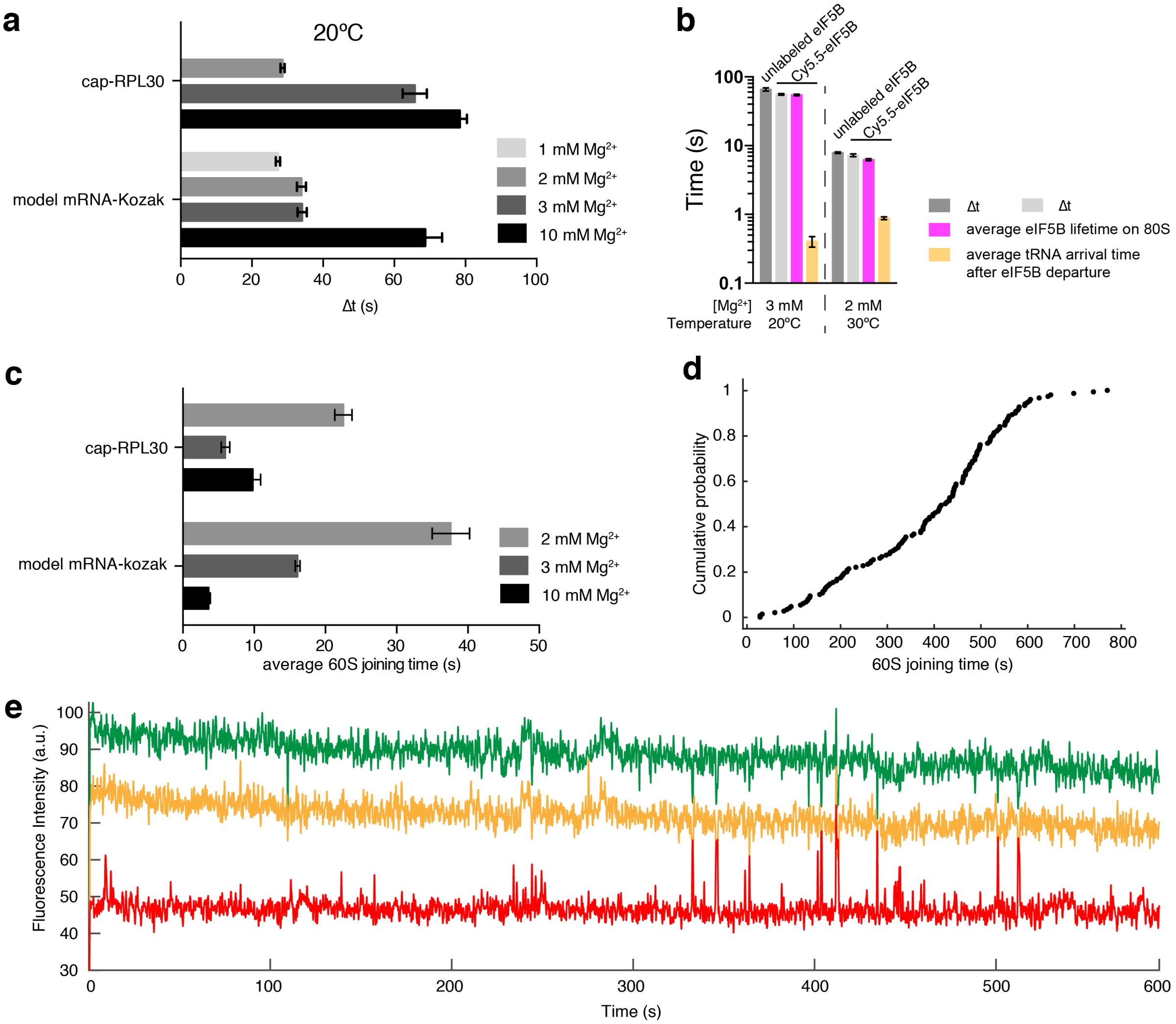


**Extended Data Fig. 7. Free $Mg^{2+}$ concentration modulates 60S joining and the transition to elongation.**
**a**, The Δt values from experiments performed with unlabeled eIF5B and cap-RPL30 mRNA or model mRNA-Kozak at 20ºC in the presence of 1 to 10 mM $Mg^{2+}$ (the data for 3 mM $Mg^{2+}$ were taken from **Fig. 2d**). For cap-RPL30, unstable 80S formation was observed at 1 mM $Mg^{2+}$ (see (**e**) below) and thus no Δt value was obtained. From bottom to top for each bar, n = 108, 189, 124, 150, 144, 118 and 195.
**b**, The Δt values, average eIF5B lifetimes on 80S, and average tRNA arrival times after eIF5B departure, from experiments performed with cap-RPL30 mRNA at 3 mM free $Mg^{2+}$ and 20ºC in the presence of unlabeled (n = 195, data taken from **Fig.2d**) or Cy5.5-eIF5B (n = 164, data taken from **Fig.3d**); or at 2 mM free $Mg^{2+}$ and 30ºC in the presence of unlabeled (n = 152, data taken from **Fig. 3f**) or Cy5.5-eIF5B (n = 150)
**c**, The average 60S joining times from the same experiments as in (**a**). The 60S arrival kinetics were fit by single- (for model mRNA-Kozak) or double- (for cap-RPL30, with the fast phase average times were plotted here) exponential functions.
**d**, The kinetic curve showing compromised 60S joining rate in experiments performed with the model mRNA-Kozak at 1 mM $Mg^{2+}$ and 20ºC as in (**a**). However, we still observed that A-site tRNA arrival occurred readily after 80S formation. The kinetics under this reaction condition were not well fit by a single- nor double-exponential distributions and therefore no average time was deduced for the bar plot in (**c**).
**e**, Sample trace from experiments performed with the cap-RPL30 mRNA at 1 mM $Mg^{2+}$ and 20ºC as described in (**a**).
Error bars in (**a**), (**b**) and (**c**) represent the 95% confidence intervals from fitting of the lifetimes to single- (**a**, **b** and **c**) or double- (**c**) exponential distributions.

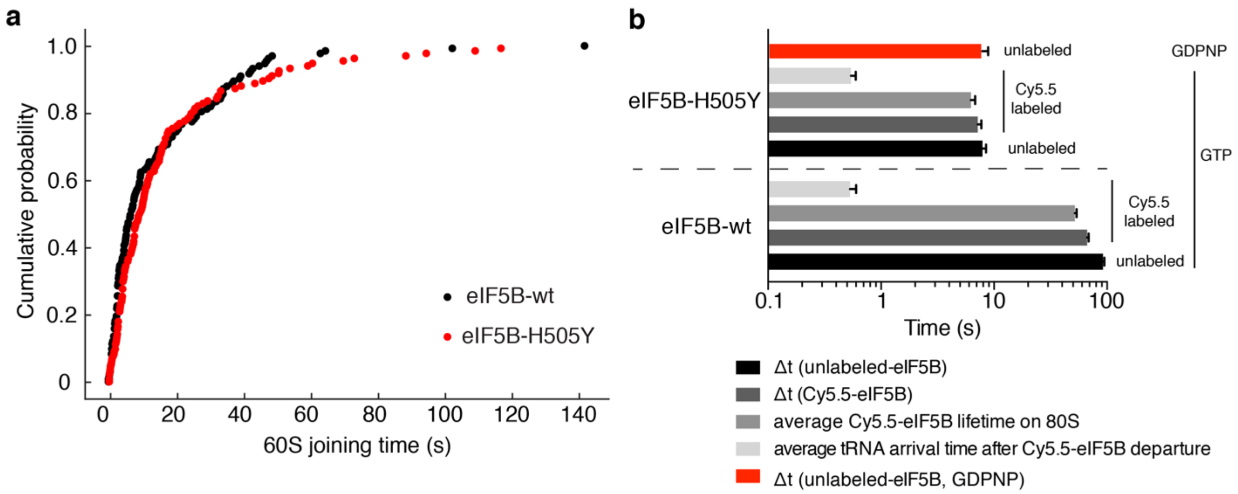


**Extended Data Fig. 8. A GTPase-defective mutant eIF5B with weaker ribosome binding affinity alters the rate of the transition to elongation.**
**a**, The kinetic curves of 60S joining from experiments performed with unlabeled wild-type (wt, n = 164) or His505Tyr mutant (H505Y, n = 119) eIF5B and the model mRNA at 20ºC in the presence of 3 mM $Mg^{2+}$ and 1mM GTP. The 60S joining rate was not significantly affected by the mutation in eIF5B.
**b**, The Δt values, average eIF5B lifetimes on 80S, and average tRNA arrival times after eIF5B departure, from experiments performed with the model mRNA at 3 mM free $Mg^{2+}$ and 20ºC, in the presence of GTP with either unlabeled eIF5B-wt (n = 164, data taken from **Fig. 2d**); Cy5.5-labeled eIF5B-wt (n = 134, data taken from **Fig. 3d** ); unlabeled eIF5B-H505Y (n = 119); or Cy5.5-labeled eIF5B-H505Y (n = 80); or separately in the presence of GDPNP with unlabeled eIF5B-H505Y (n = 95). Error bars represent the 95% confidence intervals from fitting of the lifetimes to single-exponential distributions.
